# Fentanyl and Alcohol Co-Exposure Induce Robust, Sustained Hyperlocomotion and Neural Circuit Disruption in Larval Zebrafish

**DOI:** 10.1101/2025.07.08.663662

**Authors:** Courtney Hillman, James Kearn, Matthew J. Winter, Matthew O. Parker

**Author notes:** Declaration of interest: none. Correspondence concerning this article should be addressed to Matthew O. Parker, PhD. Surrey Sleep Research Centre, University of Surrey, Guildford, GU2 7XH, UK. Email addresses of authors: Courtney Hillman, BSc., James Kearn, PhD., Matthew J. Winter, PhD, Matthew O. Parker, PhD.

## Abstract

The constantly evolving trends in substance abuse are a major concern for health authorities worldwide, and these trends include an increasing prevalence of poly-substance misuse. For instance, opioid overdoses are frequently accompanied by alcohol co-use, yet the combined effects of these substances remain poorly understood. Using zebrafish as a highly relevant vertebrate model of neuropharmacology, we investigated the interactions between fentanyl and alcohol, uncovering a unique and robust hyperlocomotor response characterized by initial locomotor suppression followed by persistent, erratic hyperlocomotion. Alcohol was found to be critical to this phenomenon, as substitution with other GABA_A_ modulators failed to replicate the effect. However, the response was replicable with heroin and remifentanil, suggesting an opioid-class wide effect. *In vivo* whole brain imaging further demonstrated dysregulated neuronal activity, with co-administration with alcohol causing potentiated neuronal activity compared to individual drug exposures and controls. Collectively, these findings suggested an integral role of ethanol and fentanyl co-administration in dysregulated neuronal responses and reveals a complex neurobehavioral mechanism. These observations suggest further investigation is warranted into the use of the larval zebrafish model for studying the neuropharmacological interactions of multiple substances of abuse.

## 1. Introduction

The growing abuse of synthetic opioids, such as fentanyl, represents a public health crisis (Arendt, 2021; CDC, 2023; Johnson et al., 2021; Lamy et al., 2021; Palamar, 2024; Rodda et al., 2017; Smith et al., 2023; Stanley, 2014; Swanson et al., 2017; United States Drug Enforcement Administration, 2022, 2024). Furthermore, the risk of overdose associated with fentanyl is especially alarming due to its high potency and increased prevalence in adulterated or misrepresented substances in the illicit drug market (Armenian et al., 2018; CDC, 2023; Rodda et al., 2017; Swanson et al., 2017; Zhu et al., 2023). Current interventions, such as treatment with the opioid receptor blocker naloxone (Narcan^®^), although lifesaving, are often less effective in cases of fentanyl overdose due to fentanyl’s rapid onset, high receptor affinity and slow clearance. In addition, the potential presence of co-ingested substances, such as alcohol, can complicate the reversal of fentanyl intoxication (Adler et al., 1997; Armenian et al., 2018; Baumann et al., 2020; Kuip et al., 2017; Leonardo et al., 2023; Montandon & Horner, 2019; Moussawi et al., 2020; Solis et al., 2018; Varshneya et al., 2019, 2021; Wagner et al., 2023; Wu et al., 2022). Recent data suggest a rising prevalence of alcohol co-abuse in individuals presenting with fentanyl overdose (Frank et al., 2015; Hickman et al., 2008; Tori et al., 2020; Witkiewitz & Vowles, 2018).

Despite these concerning trends, little is known about the pharmacokinetic and physiological interactions between fentanyl and alcohol in combination. Given that both opioids and alcohol act as central nervous system (CNS) depressants, it is probable, for example, that alcohol may amplify the risk of overdose by interfering with opioid metabolism, compounding respiratory depression, and/or complicating reversal interventions (Guerzoni et al., 2018). Indeed, evidence has shown that their co-administration can significantly increase the chance of respiratory depression, which ultimately contributes to an increased risk of overdose and fatality (White & Irvine, 1999). In addition to the risk of fatal overdose, functional interactions between the endogenous opioid system and alcohol consumption are well-documented, with evidence suggesting that the rewarding and reinforcing effects of alcohol are mediated by opioid peptide release in specific brain regions (Gianoulakis, 2001). This raises the question of how recreational co-administration of opioids and alcohol may collectively impact brain neurobiology and behavioral responses in both the short and long term.

Despite the prevalence of co-use, the neurobehavioral effects of fentanyl and alcohol remain poorly understood, and their potential for compounding harm is often underestimated (Tori et al., 2020). To address this data gap, we employed 4 days post-fertilization (dpf) larval zebrafish (*Danio rerio*) in a validated light/dark behavioral assay (Hillman et al., 2024b, 2024a), alongside *in vivo* Ca^2+^-transient mediated whole brain functional imaging, to investigate the mechanisms by which fentanyl and alcohol (ethanol) interact to produce unexpected and potentially hazardous outcomes. Integrating these approaches, we reveal the neural consequences of co-exposure of fentanyl and alcohol and provide insights to inform treatment strategies for elicit poly-substance use.

## 2. Materials and Methods

### 2.1. Animal Husbandry and Ethics

Adult and larval zebrafish (wild-type AB strain and transgenic *elavl3:GCaMP6*) were reared under standard conditions (see **Supplementary Materials: Materials and Methods** for further information). Briefly, adult animals were held at 25–28°C under standard diurnal lighting cycles and fed on a mixed diet of dry and live food. Larvae were collected after pair-wise spawning of adults and raised at circa 28°C under the same conditions until 4 dpf, when they were used in experiments. Following behavioral or imaging-based assessment, all larvae were euthanized with an overdose of anaesthetic following ethical guidelines approved by the Animal Welfare and Ethical Review Boards (AWERBs) at the Universities of Surrey and Exeter. Adult behavioral responses to fentanyl-ethanol were carried out under licence from the UK Home Office (PP8708123).

### 2.2. Chemical Exposures

Larvae were exposed to each test compound, individually or in combination, at concentrations determined through prior laboratory studies, alongside vehicle controls (0.5% DMSO), except for ethanol co-administration studies to avoid a double solvent effect. For ethanol co-administration studies, fentanyl was dissolved directly in ethanol to avoid the requirement for DMSO as an additional solvent. For complete details of the administration, see **Supplementary Materials: Materials and Methods**. The various test chemicals applied are summarized in **Supplementary Table 1**). Stock solutions of all chemicals were prepared in 100% DMSO (except for ethanol co-administration to avoid double solvent effects) and diluted to working concentrations in embryo medium immediately before use.

### 2.3. Behavioral Assays

Larval behavior was assessed using light/dark and light-only assays in Zantiks MWP units (Zantiks Ltd., Cambridge, UK), which allowed the automated tracking of locomotor activity in multiple larvae simultaneously. In the light/dark assay, larvae were subjected to alternating 5-minute phases of white light (350 lx) and darkness, with behavior being tracked for 0-30-minute post-drug exposure. Following this, a 30-minute untracked ‘break’ under light only conditions was applied to prevent acclimation to the light/dark conditions (Hillman et al., 2024a). This was then followed by a second light/dark period with recording at 60-90-minutes post-drug exposure. The light-only assay assessed locomotor responses under continuous white light for up to 80-minutes. Baseline activity was recorded prior to drug exposure, and all behavioral responses were normalized to baseline values. Adult behavior was assessed for 10-minutes in the light (350 lx) in an open-field locomotor assay using Zantiks AD units. Detailed behavioral protocols and analysis pipelines are outlined in the **Supplementary Materials: Materials and Methods**.

### 2.4. *in vivo* Whole-Brain Imaging

To investigate neuronal activity, 4 dpf transgenic *elavl3:GCaMP6* larvae were imaged using a custom-built light sheet microscope following the method of Winter *et al*. 2017 (see **Supplementary Materials: Materials and Methods** for full details). Briefly, larvae were pre-treated for 20-minutes with either 0.5% (v/v) DMSO, 1 μM fentanyl (0.5% v/v DMSO), 2.0% ethanol or a 1 μM fentanyl and 2.0% ethanol combination, following which they were immobilized using a neuromuscular blocker (4 mM tubocurarine), and embedded in low-melting point agarose within borosilicate capillaries. Fluorescence images were captured repeatedly across 10 Z-planes over 200 cycles to visualize whole-brain activity over approximately a 6-minute period. Data were processed using a custom Python pipeline, with registration to a reference brain (Randlett et al., 2015) to localize regions of interest (ROIs) in 3D.

### 2.5. Statistical Analyses

Behavioral data were analyzed using a zero-inflated gamma generalized linear mixed model (ZIG-GLMM) to address skewed movement distributions. Outliers were excluded using the median absolute deviation method. Locomotor phenotypes, including darting and steady swimming, were assessed against vehicle controls using either Kruskal-Wallis or a one-way analysis of variance (ANOVA) and pairwise post-hoc tests. Neuronal activity data were compared between exposure groups using t-tests and visualized through fluorescence temporal profiles. A full statistical breakdown is provided in the **Supplementary Materials: Materials and Methods**.

### 2.6. Whole-Brain Imaging Analyses

Image data were analyzed using a custom Python pipeline with Scikit-image (van der Walt et al., 2014), SciPy (Virtanen et al., 2020), and Scikit-Learn (Pedregosa et al., 2011). Fluorescence data were baseline-corrected and aligned to the Z-brain Atlas using affine transformations. Median fluorescence intensity within 45 brain regions were compared between exposure groups and controls using t-tests following normality checks. Significant peaks were identified based on a two-standard-deviation threshold, with key metrics (peak height, width, interval, and AUC) extracted. Data were visualized using GraphPad Prism (GraphPad Software, 2024), Excel (Microsoft Corporation, 2024), Python (Rossum & Drake, 2009) and R (R Core Team, 2024). Full analysis details are provided in **Supplementary Materials: Materials and Methods** and codes are available on GitLab and OSF.

## 3. Results

### 3.1. Co-Administration of Fentanyl and Alcohol Induces Robust Hyperlocomotor Responses in Larval Zebrafish

Fentanyl was selected for study as globally it is one of the most abused opioids and is often detected as an adulterant in illicit street drugs (Arendt, 2021; Han et al., 2019; Palamar, 2024; Rodda et al., 2017; United States Drug Enforcement Administration, 2024; Wagner et al., 2023). Alcohol was also chosen due to its status as the most abused substance and because it is frequently present in toxicological screenings of individuals experiencing fatal opioid overdose (Molina et al., 2014; Morentin et al., 2023; Winger et al., 1983). Previous work in our laboratory has established the independent concentration-dependent effects of fentanyl (Hillman et al., 2025) and ethanol (Hillman et al., 2024a) on 4 dpf larval zebrafish behavior using the light/dark assay. The light/dark paradigm introduces abrupt environmental changes that evoke robust arousal and startle responses, providing an opportunity to investigate how fentanyl-ethanol co-administration disrupts the normal behavioral responses to light and dark phase transitions. First, using a range of previously characterized concentrations in 4 dpf larval zebrafish (**see Supplementary Figure 1**), we identified that the combination of 1 µM fentanyl and 2.0% ethanol produced a hyperlocomotor phenotype in the light/dark locomotor assay.

Control larvae (water-exposure only) displayed the typical light phase hypolocomotion and dark phase hyperlocomotion. When administered independently, 2.0% ethanol induced transient changes in locomotion, with an initial decrease in dark phase locomotion, followed by a gradual increase by the final dark phase (**Figure 1A**). A similar response was seen with 1 μM fentanyl but with a full return to control-level locomotion by the final dark phase (time-dependent dark phase GLMM with gamma distribution: β = 0.023, SE = 0.0016, z = 14.79; LRT: χ²(1) = 211.86, *p* < 0.001). Following co-administration, fentanyl-ethanol exposure resulted in a highly time-dependent response (*p* < 0.001; **Figure 1A)**, characterized by a complete suppression of movement in the first dark phase, followed by a gradual increase in the second dark phase, through to robust hyperlocomotion throughout the final dark phase (from 25-minutes post-exposure). Further analysis revealed significant differences in dark phase total movement (*p* = 0.032), zero movements (*p* = 0.015), steady swim movement (*p* < 0.0001) and slow swim movement relative to control (*p* < 0.0001; **Figure 1C and E**).

**Figure 1:**
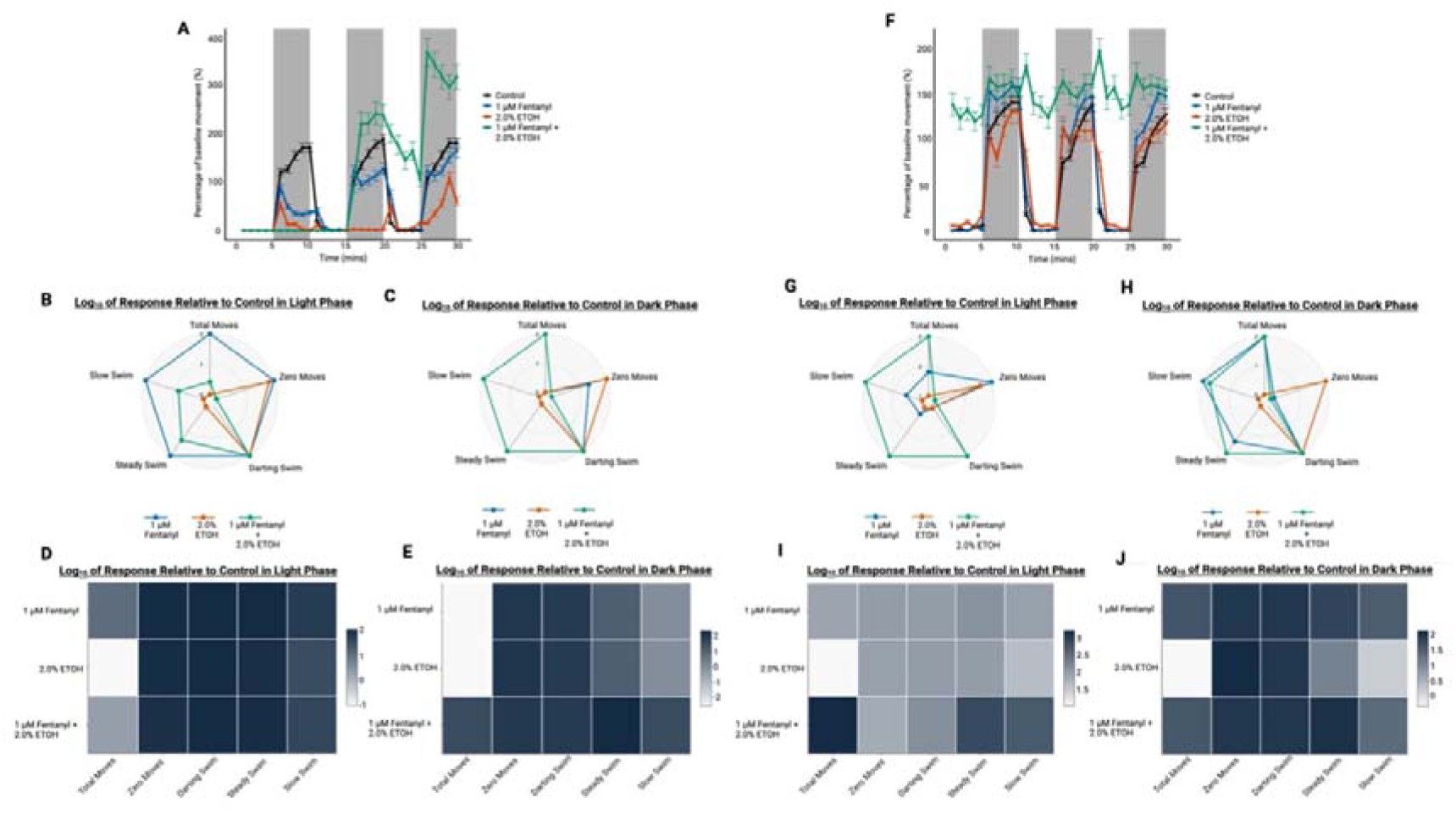
The light/dark behavioural responses to 1 μM fentanyl and 2.0% ethanol co-administration in 4 dpf zebrafish larvae. The light/dark minute analysis for 0-30-minutes post-exposure (**A**) and 60-90-minutes post-exposure (**F**) represented as mean percentage of baseline movement. Light phase swim phenotype responses shown as a radar chart and heat map in the 0-30-minute recording (**B** and **D**) and the 60-90-minute recording (**G** and **I**). Dark phase swim phenotype responses are shown as radar chart and heat map in 0-30-minute recording (**C** and **E**) and the 60-90-minute recording (**H** and **J**). Data in A and F are presented as mean ± standard error of the mean (SEM). Data in **B – E** and **G – J** are presented as mean. (*n* = 98, observed power = 0.997).

After one hour of exposure, the observed effect of individual fentanyl and ethanol treatment alone had returned to control light/dark activity levels (**Figure 1F; see Supplementary Video 1** for visual representation of the behavioral response). However, the fentanyl-ethanol combination resulted in persistent hyperlocomotion across both light and dark phases, with the activity level at each point being significantly higher than corresponding vehicle-controls (*p* < 0.001). Although the distinct light/dark phase response was almost fully overshadowed by the hyperlocomotion, there was some evidence of distinct increases in activity within the first minute of transition from dark to light phase in the 2^nd^ and 3^rd^ light phases, suggesting a possible sensory-linked arousal component to the hyperlocomotor response.

To investigate if the hyperlocomotion represented a sensory-linked arousal component, we analyzed the difference in locomotion between the minute before, minute of and minute post the light phase changes (**Supplementary Figure 2**). However, no significant difference was seen for the pre-, during and post-light phase changes suggesting movement with fentanyl-ethanol co-exposure is largely independent of sensory stimuli.

Further analysis of larval locomotor patterns at 60-90 minutes exposure showed significantly increased total movements in the dark phase (*p* = 0.0007) and increased in the light (*p* < 0.0001). A significant increase in steady swim responses (*p* < 0.0001) and slow swim responses in the light phase (*p* < 0.0001) were also reported with the combined exposure compared to controls (Hillman et al., 2024a). Interestingly, no significant difference was seen in the dark for darting moves (*p* = 1.0), but a significant difference was seen in the light phase (*p* < 0.0001).

The observed locomotor effects reveal significant and distinctive behavioral changes, particularly in swim phenotypes, which are not typical of single-substance exposures (Hillman et al., 2024a, 2025) (see **Supplementary Materials: Swim Phenotypes** for more information). These findings suggest a possible dual mechanism involving both arousal and sensory pathways. Initially the combination of fentanyl and ethanol appears to suppress locomotion, followed by a pronounced increase in activity, independent of the light conditions.

### 3.2. The Observed Hyperlocomotor Phenotype is Replicated in Adult Zebrafish

We next confirmed the behavioral phenotype was not a larval-specific response by looking to see whether the observed hyperlocomotion could be replicated in adult zebrafish using an open-tank behavioral assay and immersion exposure (**Supplementary Figure 3 and Supplementary Video 2**). For the adult fish, light/dark stimulation was not used, as this response is only relevant in larvae (Hillman et al., 2024a). For comparison with the adult data, the larval light-only responses can be seen in **Supplementary Figure 4**. We used 1.0% ethanol exposure in adults (rather than 2%) following extensive previous work with ethanol in our laboratory, preliminary concentration-response assessment, and with data from other researchers that have shown 2.0% ethanol to be toxic in adults (Lin et al., 2015). We found that the larval hyperlocomotor phenotype can broadly be replicated in adult zebrafish exposed to a combination of 1 μM fentanyl and 1.0% ethanol compared to controls (*p* = 0.013). Adults do not exhibit the initial pronounced suppression of locomotion that is seen in larvae, highlighting some age-related differences.

### 3.3. Substitution Experiments Highlight Opioid-Specific Mechanisms

To evaluate the specificity of the fentanyl response, we substituted fentanyl with remifentanil and diacetylmorphine (heroin) (**Figures 2 and 3**), as two alternative opioid receptor agonists.

**Figure 2:**
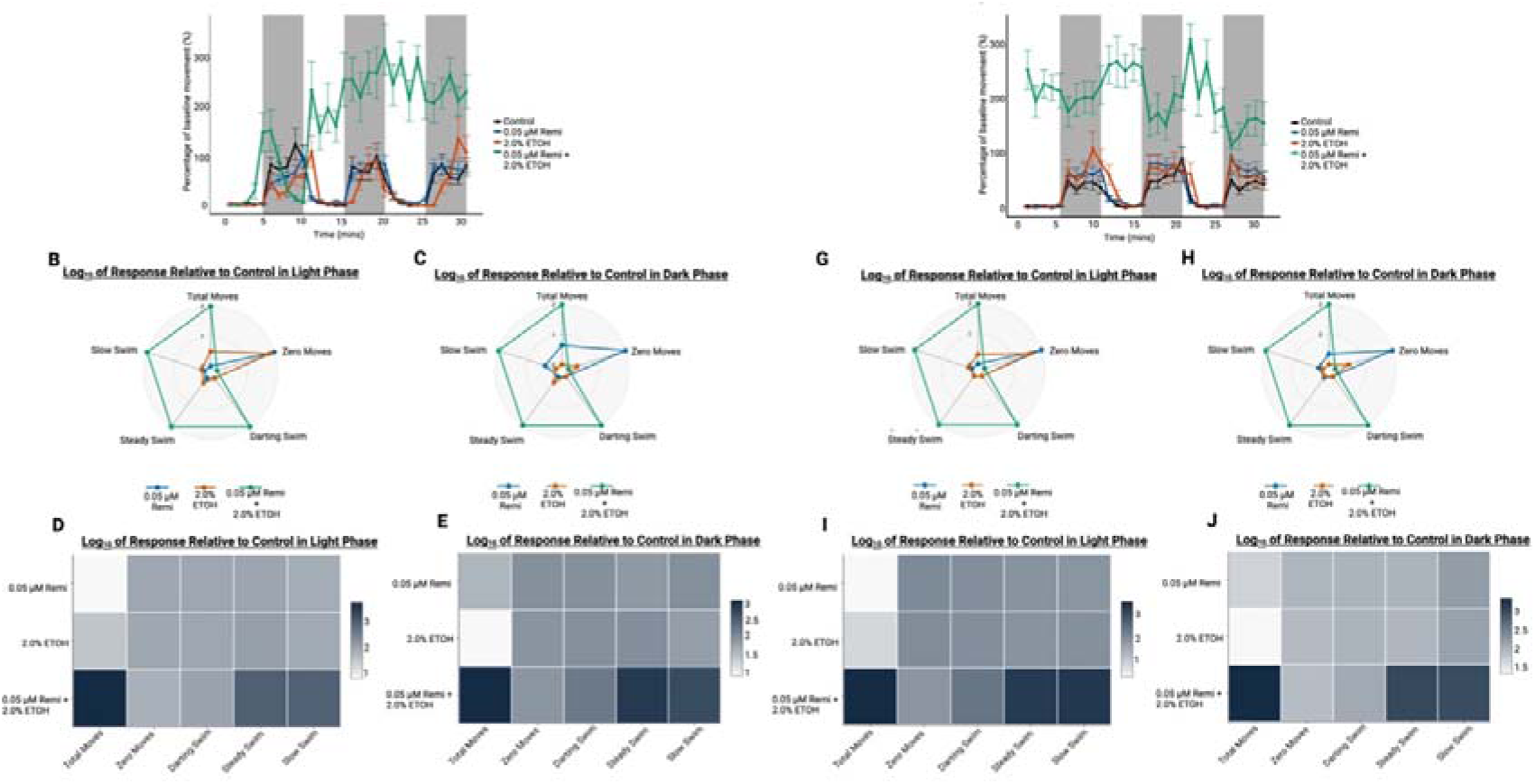
The light/dark behavioural responses to 0.05 μM remifentanil (Remi) and 2.0% ethanol co-administration in 4 dpf zebrafish larvae. The light/dark minute analysis for 0-30-minutes post-exposure (**A**) and 60-90-minutes post-exposure (**F**) represented as mean percentage of baseline movement. Light phase swim phenotype responses shown as a radar chart and heat map in the 0-30-minute recording (**B** and **D**) and the 60-90-minute recording (**G** and **I**). Dark phase swim phenotype responses are shown as radar chart and heat map in 0-30-minute recording (**C** and **E**) and the 60-90-minute recording (**H** and **J**). Data in A and F are presented as mean ± SEM. Data in **B – E** and **G – J** are presented as mean. (*n* = 16, observed power = 0.971).

**Figure 3:**
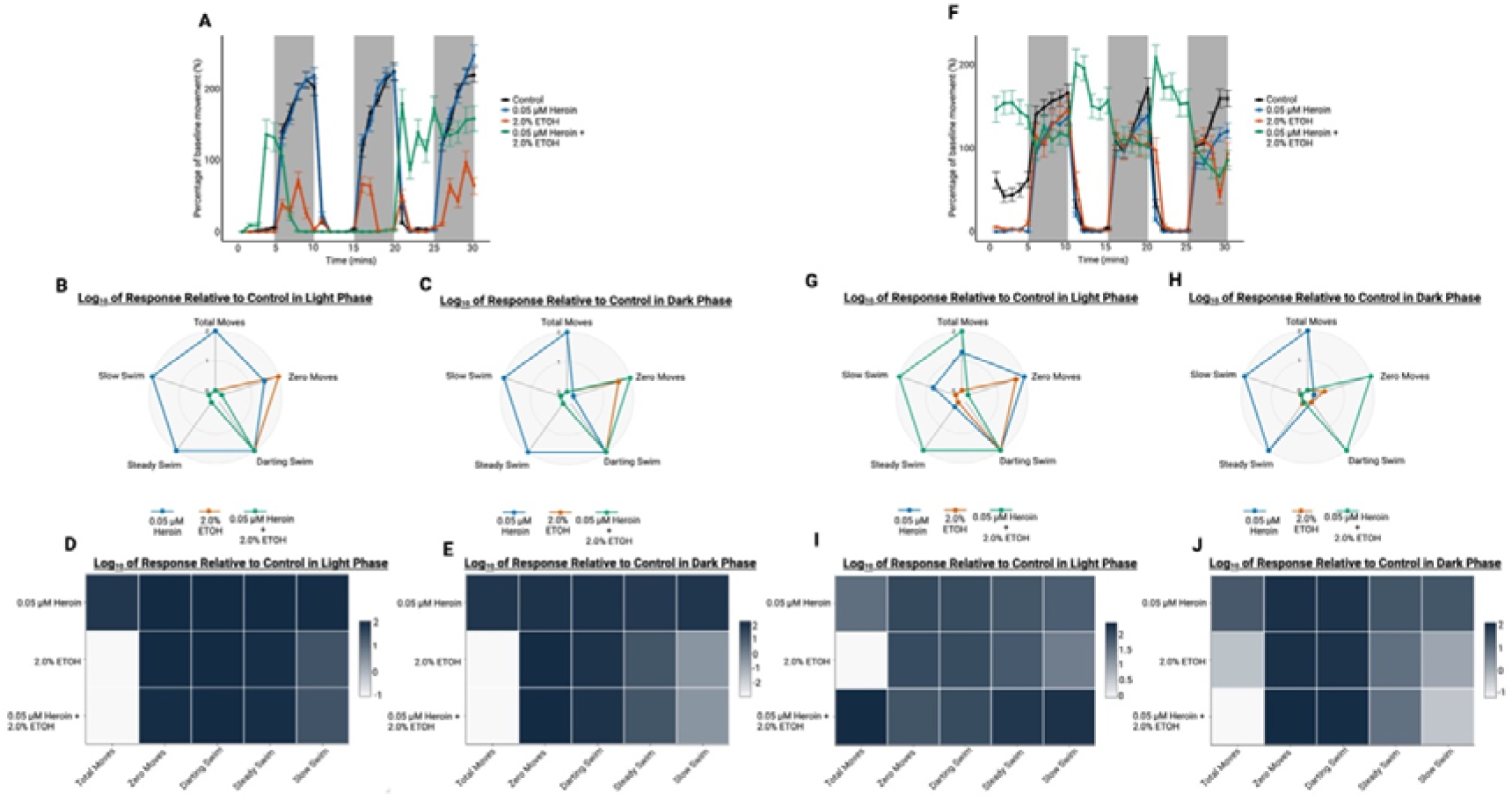
The light/dark behavioural responses to 0.05 μM heroin and 2.0% ethanol co-administration in 4 dpf zebrafish larvae. The light/dark minute analysis for 0-30-minutes post-exposure (**A**) and 60-90-minutes post-exposure (**F**) represented as mean percentage of baseline movement. Light phase swim phenotype responses shown as a radar chart and heat map in the 0-30-minute recording (**B** and **D**) and the 60-90-minute recording (**G** and **I**). Dark phase swim phenotype responses are shown as radar chart and heat map in 0-30-minute recording (**C** and **E**) and the 60-90-minute recording (**H** and **J**). Data in A and F are presented as mean ± SEM. Data in **B – E** and **G – J** are presented as mean. (*n* = 63).

The combination of remifentanil-ethanol triggered a faster (compared to fentanyl-ethanol) and more erratic increase in locomotion, detectable within 10 minutes post-exposure and persisting through the 60-90-minute period (**Figure 2A**). This hyperlocomotion occurred across both light and dark phases (GLMM: β = 0.83, SE = 0.07, z = 11.66; LRT: χ²(1) = 124.78, *p* < 0.001). Notably, remifentanil-ethanol elicited unique swim phenotype patterns compared to remifentanil and ethanol independent exposures, including significantly increased total movements (*p* < 0.0001 [light], *p* < 0.0001 [dark]), darting moves (*p* = 0.0038 [light], *p* < 0.0001 [dark]), steady swim (*p* < 0.0001 [light], *p* < 0.0001 [dark]) and slow swim (*p* < 0.0001 [light], *p* < 0.0001 [dark]) moves during the 0-30-minute period. By the 60-90-minute period, hyperlocomotion was observed across both phases, with significant increases in total movements (*p* = 0.002 [light], *p* < 0.0001 [dark]), darting moves (*p* < 0.0001 [light], *p* = 0.011 [dark]), steady swim moves (*p* < 0.0001 [light], *p* < 0.0001 [dark]) and slow swim (*p* < 0.0001 [light], *p* < 0.0001 [dark]). A significant increase in zero movements was also seen in the light phase only compared to vehicle-controls (*p* < 0.0001).

Heroin-ethanol co-exposure resulted in comparable co-exposure responses to the fentanyl-ethanol response (**Figure 3**). During the first 30 minutes post-exposure, heroin-ethanol significantly decreased total movements (*p* < 0.0001 [light], *p* < 0.0001 [dark]), steady swim (*p* = 0.0252 [light], *p* < 0.0001 [dark]) and slow swim (*p* < 0.0001 [light], *p* < 0.0001 [dark]) responses (**Figure 3**). Unlike fentanyl- or remifentanil-ethanol combinations, however, heroin-ethanol produced a distinctive profile at 60-90 minutes, with significantly decreased dark phase total movements (*p* < 0.0001), steady swim moves (*p* < 0.0001), and slow swim moves (*p* < 0.0001) with significantly increased darting moves (*p* < 0.0001) compared to controls. Whereas, a significant increase in light phase total moves (*p* < 0.0001), steady swim (*p* < 0.0001) and slow swim (*p* < 0.0001) was seen. These swim phenotype responses match the behavioral pattern seen, with a suggested reversal of light/dark responses with heroin-ethanol co-exposure characterised by a greater degree of light phase responsiveness seen compared to dark phase locomotion, which was not seen with fentanyl- or remifentanil-ethanol combinations. In addition, the heroin-ethanol co-exposure saw a transient increase in locomotion at around 3-7 minutes prior to the onset of the total suppression of locomotion that precedes the sustained hyperlocomotion, a response not seen with the fentanyl-ethanol and remifentanil-ethanol exposure. These results highlight key differences in behavioral responses depending on the opioid combined with ethanol. However, despite these differences, the overall effect of combining ethanol with opioids is sustained hyperlocomotion, suggesting a common physiological interaction across the opioid class.

### 3.4. Ethanol substitution with the **γ**-aminobutyric acid (GABA)_A_ positive allosteric modulator, diazepam, reveals ethanol specificity

Ethanol exerts its effects through numerous neurotransmitter systems with predominant behavioral effects mediated by GABA_A_ receptor positive allosteric modulation (Lobo & Harris, 2008). Therefore, to probe the role of ethanol in the co-exposure response we substituted it with diazepam, a positive allosteric modulator (PAM) of the GABA_A_ receptor (Campo[Soria et al., 2006). Exposure to the fentanyl-diazepam combination significantly reduced dark phase locomotion during both the 0-30-minute and 60-90-minute post-exposure periods compared to vehicle controls (*p* < 0.001 for both; **Figure 4A, F**). This decrease was also observed across both light and dark phases when comparing the fentanyl-ethanol and fentanyl-diazepam combinations during the 60-90-minute period (*p* < 0.001). Additionally, diazepam-fentanyl exposure altered swim phenotypes, including decreased total moves (*p* < 0.0001 [light], *p* < 0.0001 [dark]), zero moves (*p* < 0.0001 [dark]), steady swim (*p* < 0.0001 [light], *p* < 0.0001 [dark]) and slow swim (*p* < 0.0001 [dark]) compared to controls in the 0-30-minute period. Increased slow swim was also seen in the dark phase (*p* < 0.0001). However, by the 60-90-minute period an increase in total moves is seen in the dark phase (*p* < 0.0001) with increased slow swim predominantly seen for both phase (*p* < 0.0001, for both). Despite the increased dark phase total moves, we also report significantly increased zero moves with the fentanyl-diazepam exposure compared to controls (*p* = 0.0001). Importantly, no hyperlocomotion was seen with the fentanyl-diazepam co-exposure as is seen with fentanyl-ethanol, confirming the hyperlocomotor response is specific to ethanol rather than generic GABAergic modulation. This suggests that ethanol potentiates fentanyl’s behavioral responses during co-exposure.

**Figure 4:**
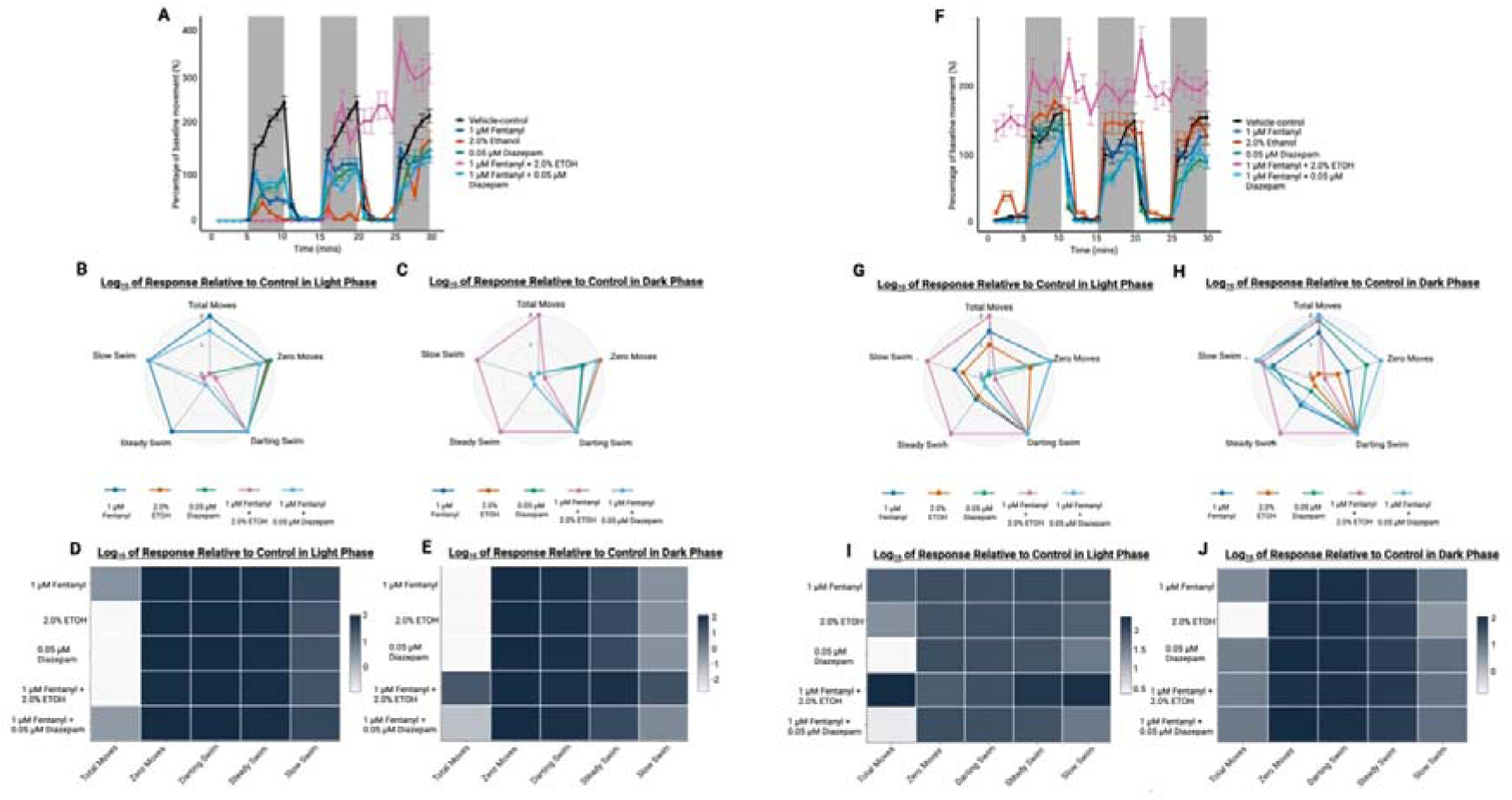
The light/dark behavioural responses comparing the fentanyl-ethanol combination to fentanyl co-administered with 0.05 μM diazepam, in 4 dpf zebrafish larvae. The light/dark minute analysis for 0-30-minutes post-exposure (**A**) and 60-90-minutes post-exposure (**F**) represented as mean percentage of baseline movement. Light phase swim phenotype responses shown as a radar chart and heat map in the 0-30-minute recording (**B** and **D**) and the 60-90-minute recording (**G** and **I**). Dark phase swim phenotype responses are shown as radar chart and heat map in 0-30-minute recording (**C** and **E**) and the 60-90-minute recording (**H** and **J**). Data in A and F are presented as mean ± SEM. Data in **B – E** and **G – J** are presented as mean. (*n* = 76).

### 3.5. Dysregulated Brain Activity with Evidence of Complex Neurobehavioral Mechanisms

To investigate the impact of fentanyl-ethanol co-exposure directly on brain function we employed *in vivo* whole brain functional imaging, exploiting the optical transparency of transgenic 4 dpf larvae expressing the genetically encoded Ca^2+^ indicator *GCaMP6* under control of the pan-neuronal *elavl3* promoter (*Tg*:(*elavl3:GCaMP6s*)) (Chen et al., 2013; Vladimirov et al., 2014). This imaging-based method allows live visualization of neural activity across intact brain structures with or without the presence of drug treatment (Winter *et al*., 2021).

The results of the functional imaging revealed that the fentanyl-ethanol temporal profile differs substantially from the individual drug exposures and controls, with broadly elevated GCaMP fluorescence indicating widespread activation of neuronal populations. Exposure to fentanyl alone showed a modest increase in GCaMP peak amplitude and decrease in the number of peaks across most brain regions compared with controls, a pattern that was greatly exacerbated when both drugs are co-administered (**Figure 5**). Specifically, at approximately 100 seconds of recording (∼25-mins post-exposure) a complete loss of distinct peaks is seen with a pronounced increase in GCaMP fluorescence regardless across all brain regions. This is mirrored in the decreased number of peaks (**Figure 6A**) with the combination exposure compared to controls and ethanol (*p* < 0.0001, respectively) and an increase in the average peak amplitude (**Figure 6B**) with the fentanyl-ethanol combination compared to controls (*p* = 0.01) and ethanol (*p* < 0.0001). The variability in peak number and peak amplitude in the fentanyl-ethanol combination was greater than that seen with the controls and singly drug-exposed larvae, which may in part be due to the biphasic nature of the fentanyl-ethanol response.

**Figure 5:**
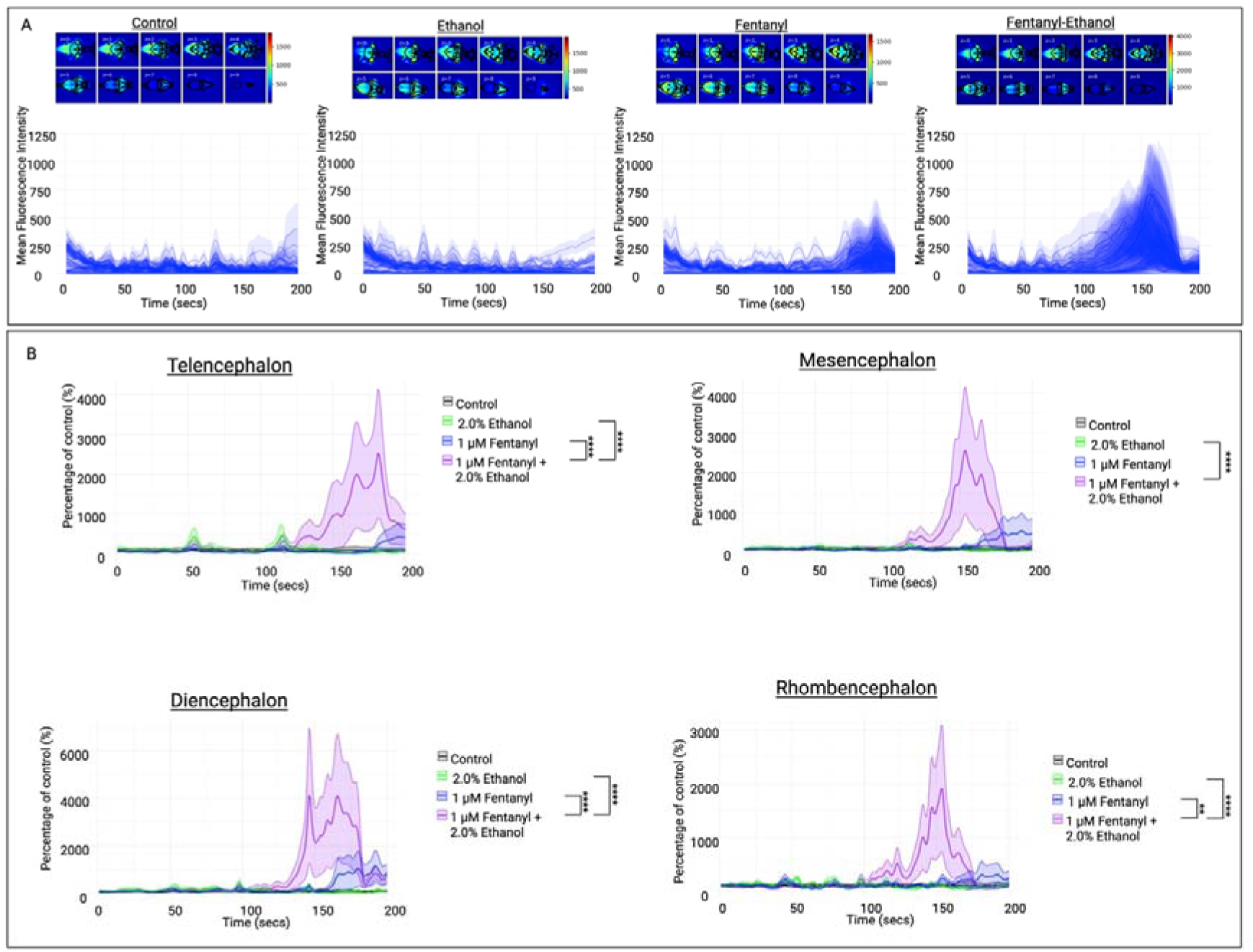
Temporal profiling and median florescence projections following functional 4-dimensional brain profiling in fentanyl-ethanol treated 4 dpf zebrafish larvae. (**A**) The median fluorescence intensity projections across time for each z-plane (0-9 ventral to dorsal surface) and corresponding temporal profile with each line representing the median intensity of the voxels within each of the 3D-registered anatomical regions for controls, 2.0% ethanol, 1 μM fentanyl and fentanyl-ethanol combination exposures. (**B**) the temporal profiles of GCaMP fluorescence after ethanol, fentanyl and fentanyl-ethanol exposure as a percentage of control fluorescence in the telencephalon, diencephalon, mesencephalon and rhombencephalon. *(n = 7 larvae). Significance values are set as: **** p < 0.0001, *** p < 0.001, ** p < 0.01, * p < 0.05*.

**Figure 6:**
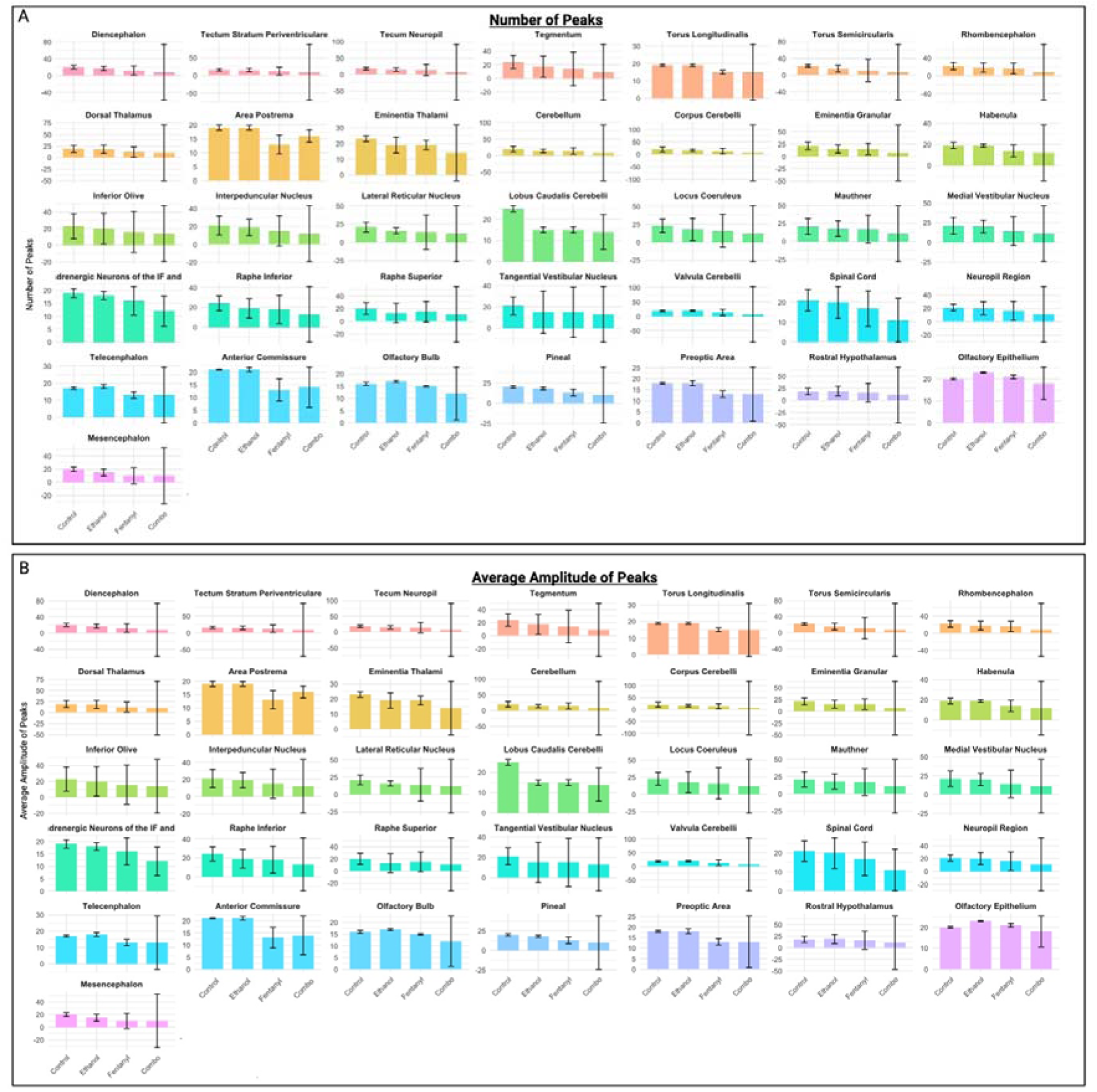
The number of fluorescence peaks and amplitude following functional 4-dimensional brain profiling in fentanyl-ethanol treated 4 dpf zebrafish larvae. **(A**) The mean number of peaks in GCaMP fluorescence in each brain region measured in the control, ethanol, fentanyl and the fentanyl-ethanol combination exposed animals. (**B**) The average amplitude of GCaMP fluorescence peaks in brain regions from the control, ethanol, fentanyl and fentanyl-ethanol exposed larvae. *(n = 7 larvae). Significance values are set as: **** p < 0.0001, *** p < 0.001, ** p < 0.01, * p < 0.05*.

These observations, coupled with the trend in increased forebrain fluorescence seen compared to individual fentanyl and ethanol exposures (**Supplementary Figure 5**), resulted in our further investigation into the neuroanatomical differences in neuronal activity observed. We revealed an apparent potentiation of fentanyl neuronal responses when combined with ethanol regardless of brain region (**Figure 5B**). In both the telencephalon and diencephalon, a significant increase in fluorescence is seen with the fentanyl-ethanol combination compared to both fentanyl and ethanol (*p* < 0.0001). This was also reported in the rhombencephalon; however, with a substantially reduced mean GCaMP fluorescence compared to the diencephalon and telencephalon. Comparatively, in the mesencephalon, only the ethanol exposure was significantly different to the fentanyl-ethanol combination (*p* < 0.0001). These traces also highlight other areas of potential future investigation; for instance, at approximately 150 seconds post-exposure we see an increase in fluorescence responses with fentanyl-alone for all four anatomical regions. This suggests a potential time-dependent neuronal response to fentanyl, which may play a role in the extended behavioral response we see with the fentanyl-ethanol combination. However, this requires further investigation to confirm.

## 4. Discussion

Toxicology screening often detects ethanol in individuals presenting with opioid overdose, however the physiological and neurobehavioral impact of co-administration remains understudied, and the interactive pharmacological processes poorly understood (Guerrieri et al., 2017; Tori et al., 2020). We revealed a a robust response characterised by initial hypolocomotion followed by erratic and sustained hyperlocomotion followed by erratic, sustained locomotion. Further investigation revealed the response to be broadly replicable regardless of opioid used (reproduced with remifentanil and heroin). The response was not seen with fentanyl-diazepam co-exposure, suggesting it is ethanol-specific and is not solely due to GABA_A_ positive allosteric modulation. We furthered our investigation by performing neuronal analyses and confirmed extensive neuronal functional dysregulation particularly when both drugs were administered in combination. Taken together, our findings highlight numerous potential avenues for future research into the pharmacological and neuronal mechanisms underlying this response.

We confirmed the response to be specific to ethanol by substituting ethanol with diazepam. Diazepam is a PAM of the GABA_A_ receptor (Campo[Soria et al., 2006) and GABAergic transmission is highly sensitive to alcohol, with GABA_A_ receptor antagonists effectively blocking the reinforcing effects of alcohol (Cruz et al., 2008). In addition to the known reinforcing effects attributed with GABAergic neurotransmission, alcohol is also known to enhance GABA_A_ function through direct receptor binding (Davies, 2003). Benzodiazepines, including diazepam, are known to mimic alcohol intoxication effects including anxiolysis, sedation and impaired motor function, thought to be through GABA_A_ receptor effects (Goldschen-Ohm, 2022). We, therefore, postulated that the hyperlocomotion may be attributed with GABA_A_ receptor activation and tested this by substituting ethanol with diazepam. However, we found that no significant difference to fentanyl-ethanol with no hyperlocomotion seen. This may be attributed to a need for direct GABA_A_ receptor activation rather than allosteric enhancement (Campo[Soria et al., 2006). In addition to evaluating the GABA_A_ role in the response, we assessed the specificity of the opioid-driven effect by replacing fentanyl with heroin and remifentanil, both of which also resulted in sustained hyperlocomotion in combination with ethanol. Of significant interest is the prolonged and potentiated response we observed specifically with remifentanil exposure. We previously revealed a rapid return to control locomotion in our larval studies with remifentanil exposure alone (Hillman et al., 2025), which we theorised was a result of remifentanil’s unique metabolic pathway (Cascone et al., 2013). As such, the prolonged and potentiated effect with remifentanil-ethanol does not fit with our previous theories regarding remifentanil in 4 dpf larvae and therefore future research into the opioid-specific role of the response is required.

We next undertook whole brain functional imaging with an aim of ruling out the potential seizuregenic activity of the combination exposure, as well as to provide further insight into the neuronal mechanisms underlying the behavioral response. Our findings revealed a dysregulated temporal profile of neuronal activity that corresponds with the behavioral phenotype seen, characterized by substantial increases in GCaMP fluorescence indicative of widespread neuronal excitation, coupled with a loss of distinctive (control) activity. Specifically, we report a reduction in the number of peaks in brain activity regardless of brain region, coupled with increased peak amplitude, clearly seen at approximately 150 seconds post-exposure, which corresponds with the timing of the hyperlocomotion in the behavioural findings. Further analysis revealed this dysregulated fluorescence activity profile across the four distinct major neuroanatomical regions (telencephalon, diencephalon, mesencephalon and rhombencephalon), suggesting a non-specific neuronal activation with the fentanyl-ethanol combination regardless of neuroanatomical region. Although our neuronal profile does not broadly correspond with traditional fluorescence activity profiles relating to exposure to seizuregenic compounds in larval zebrafish (Turrini et al., 2022; Winter et al., 2021), we cannot rule out the response is seizure-like in nature. Similarly, our behavioral findings do not correspond with the rapid darting and whirlpooling locomotion traditionally seen with zebrafish experiencing seizures (Baraban et al., 2005; Turrini et al., 2022), yet there remains a possibility of a seizure-like response, as individual seizurogenic compounds are known to produce distinctive behavioral and electrographic responses depending on the mechanism of action (Pinion et al., 2022; Winter et al., 2021; Yang et al., 2017). Interestingly, fentanyl-alone resulted in increased neuronal fluorescence activity at approximately 150-seconds in all four neuroanatomical regions and when in combination with ethanol, this response was substantially greater. This suggested a potential role of ethanol in potentiating fentanyl responses. However, this does not correspond with our behavioral findings and as such must be interpreted with caution, requiring further investigation.

Importantly, it should be noted that the use of poly-drug exposures may have introduced unintended stress responses, potentially contributing to increases in ‘escape’-like locomotion. It is also noteworthy that the fentanyl-ethanol response is highly distinctive from that seen with single drug exposures, in particular the biphasic nature of the response, characterised by an initial complete suppression of locomotion, followed by a sustained hyperlocomotion. This stark contrast in response with time is likely to limit the effectiveness of our standardised analysis pipeline. As such, additional work may be needed to better understand both locomotion and CNS activity during the distinctive suppression and excitation phases of the fentanyl-ethanol response. Additionally, the neuronal biology and physiology of zebrafish differ from those of mammals and humans. Furthermore, A caveat of the 30-minute recording is that the time-dependent response seen (i.e., zero movement initially to hyperlocomotion) may skew our interpretation of results. Despite these potential limitations, our data provides valuable insights into conserved mechanisms, even as we acknowledge the need for further work to enhance translational applicability.

## 5. Conclusions

Our study reveals novel insights into the complex interactions between synthetic opioids and ethanol, highlighted by the fentanyl-ethanol induced hyperlocomotor response in 4 dpf larval zebrafish, with confirmation of this hyperlocomotion in adult zebrafish. We undertook substitution analyses to reveal an ethanol-specific role in the response. However, we also identified the response to be replicable across three distinct opioid drugs of abuse (fentanyl, heroin and remifentanil). We further analysed the response by incorporating neuronal calcium imaging-based approaches, which suggested the response is not seizure-like. We do however acknowledge that further research is required to confirm this. Finally, we provide further insight into the behavioral response by revealing extensive neuronal dysregulation. Therefore, we suggest future research is required to underscore the mechanisms behind the hyperlocomotion and neuronal dysregulation observed. Collectively, these findings provide novel insights into the neurobehavioral effects of opioid and ethanol co-exposure, emphasizing the potential role of ethanol in potentiating opioid effects. The insights gained may inform future studies into the potential mechanisms underlying the response and provide avenues for therapeutic intervention research for poly-substance abuse.

## 7. Declarations/Acknowledgments

### Ethics approval

All experiments were approved by the University of Portsmouth and Exeter Animal Welfare and Ethical Review Board, and under license from the UK Home Office (Animals [Scientific Procedures] Act, 1986, license number PP8708123).

### Consent to participate

Not applicable. Consent for publication: Not applicable.

### Availability of data and materials

All data and analysis materials are available on the Open Science Framework: https://osf.io/2bg9t/

### Code availability

Not applicable.

### Authors’ contributions

Conceptualization: CH, JK, MJW, MOP. Data Curation: CH. Formal Analysis: CH. Funding Acquisition: JW, MJW, MOP. Investigation: CH. Methodology: CH, JK, MJW, MOP. Project Administration: MJW, MOP. Resources: JK, MJW, MOP. Software: MJW, MOP. Supervision: JK, MJW, MOP. Validation: CH. Visualisation: CH. Writing – original draft preparation: CH. Writing – review & editing: CH, JK, MJW, MOP.

