## Supplementary for "Fentanyl and Alcohol Co-Exposure Induce Robust, Sustained Hyperlocomotion and Neural Circuit Disruption in Larval Zebrafish"

1. **Supplementary Materials**
   1.1. **Materials and Methods**
        1.1.1. **Animal Husbandry and Ethics**

The animals used in this study included wild-type (WT) and genetically modified strains bred in-house. For behavioral analysis, adult zebrafish of the AB WT strain were kept in a recirculating aquatic system (Aquaneering, Fairfield Conrolec Ltd., Grimsby, UK). Fish were housed in tanks with a 50:50 male-to-female ratio, with 8–10 fish in each 2.8 L tank and up to 20 fish in each 6 L tank. The system was maintained at 25–27°C, with a pH of 8.4 (±0.4) under a 14-hour light and 10-hour dark cycle, with lights turning on at 8:30 am. Fish were initially fed ZM-100 fry food (Zebrafish Management Ltd., Winchester, UK) from 5 dpf until adulthood, after which they were transitioned to a diet of flake food and live brine shrimp, provided three times daily on weekdays and once per day on weekends. Larvae were reared in a translucent incubator at 28.5°C prior to larval behavioral testing. Following testing, larvae and adult fish were euthanized using Aqua-Sed™ (Vetark, Winchester, UK) as per the manufacturer’s guidelines. Adult behavioral responses to fentanyl-ethanol were carried out under licence from the UK Home Office (PP8708123).

For light-sheet microscopy, transgenic zebrafish expressing the pan-neuronal calcium sensor *elavl3* were housed in recirculating aquaria at 28°C (±1°C) on a 12-hour light and 12-hour dark cycle, with a 20-minute gradual transition between light and dark phases. The recirculating water was filtered using a reverse osmosis system (Environmental Water Systems UK Ltd.) and reconstituted with analytical-grade mineral salts (117 mg/L CaCl₂, 25 mg/L NaHCO₃, 50 mg/L MgSO₄, 2.3 mg/L KCl, and 1.25 mg/L Tropic Marine Sea Salt) to maintain a conductivity of approximately 300 μS/cm. Spawning was induced by introducing a spawning substrate at dawn, and fertilized eggs were collected, rinsed, and transferred to Petri dishes containing culture water, where they were maintained until reaching 4 dpf.

All procedures were approved by the Animal Welfare and Ethical Review Boards (AWERB) at the University of Surrey and the University of Exeter and adhered to UK Home Office licensing requirements.

1.1.2. **Chemical Exposures**

**Supplementary Table 1** provides an overview of the chemicals used in the present study, including exposure concentrations. All concentrations tested were selected based on concentration-response findings in our lab.

**Supplementary Table 1:** The compounds used, their chemical class, supplier, catalogue number and the concentration tested in μM [mg/L].

| **Compound** | **Chemical Class** | **Supplier** | **Catalogue Number** | **Concentration tested [μM (mg/L)]** |
| --- | --- | --- | --- | --- |
| Diacetylmorphine (Heroin) | Opioid | Sigma-Aldrich | H159 | 0.05 [0.018] |
| Diazepam | GABA_A_ PAM | Sigma-Aldrich | D0899 | 0.05 [0.01] |
| Dimethyl Sulfoxide (DMSO) | Solvent | Sigma-Aldrich | 472301 | -- |
| Ethanol | Alcohol | Sigma-Aldrich | S8010 | -- |
| Fentanyl citrate salt (Fentanyl) | Opioid | Sigma-Aldrich | F3886 | 0.0005 – 5 [0.00026 - 2.64] |
| Remifentanil hydrochloride (Remifentanil) | Opioid | Sigma-Aldrich | Y0001892 | 0.05 [0.02] |
| Tricaine methanesulfonate (MS-222) | Anaesthetic | Sigma-Aldrich | E10521 | 7654.3 [2.0] |

All chemicals were made up as stock solutions in 100% DMSO and stored at -20°c where applicable, except fentanyl which was made up in water to avoid a double solvent effect with DMSO and ethanol. For ethanol co-administration studies, fentanyl was dissolved directly in ethanol to avoid the requirement for DMSO as an additional solvent.

1.1.3. **Behavioral Apparatus**

Behavioral testing was conducted on weekdays between 09:00 and 18:00, using Zantiks MWP units (Zantiks Ltd., Cambridge, UK). Each test was recorded from above to enable real-time observation of larval behavior within the apparatus.

1.1.4. **Light/Dark Behavioral Assay**

For the standardized light/dark assay procedure, refer to Hillman et al. (2024b) for details, with additional applications discussed in (Hillman et al. (2024a). Briefly, larvae were pre-acclimated at 28°C in a benchtop incubator located in the testing room for 30 minutes. During each test, 225 µL of water from a Petri dish, along with a randomly selected larva, was transferred into individual wells of a 48-well plate using pipette tips trimmed by about 2 mm at the tip. Larvae were selected randomly from different Petri dishes for each session. Following transfer, the well plate was placed in the Zantiks MWP unit, and overhead lights were switched on for a 30-minute acclimation period. The light/dark behavior tracking then commenced, with a 30-minute baseline period consisting of alternating five-minute phases of light (350 lx, white light) and darkness, repeated three times. Temperature control within the MWP unit was maintained precisely at 28.02°C (± 0.07°C) throughout the experiment. After the baseline recording, the well plate was removed from the unit for chemical treatment (see Chemical Exposures), applied using a multi-channel pipette, then returned to the unit for further behavior tracking under alternating light/dark conditions. To study acute effects, a 0-30-minute post-exposure recording was performed. Following this, the plate was left in the unit for a 30-minute untracked ‘break’ in light only conditions. This break was included to ensure the larvae did not acclimate to the light/dark conditions (Hillman et al., 2024a). Finally, to look at longer-term exposure, a second light/dark recording was performed at 60-90-minutes post-exposure.

1.1.5. **Light-only Behavioral Assay**

The light-only assay followed the same principles at the light/dark assay. However, instead of alternating light/dark conditions, the larvae were recorded for 80-minutes post-exposure under white light (350 lx).

1.1.6. **Adult Behavioral Assay**

Adult zebrafish (8 months post-fertilization) were immersion exposed to either water-based control, 1 μM fentanyl, 1.0% ethanol or a combination of 1 μM fentanyl and 1.0% ethanol (*n* = 4 per group (50:50 male: female)). This ethanol concentration was selected following preliminary concentration-response findings in our lab and evidence that 2.0% ethanol is toxic to adult zebrafish (Lin et al., 2015). The fish were then immediately tracked for 10-minutes in Zantiks AD behavioral units using an open field lights-on behavioral paradigm. Data were extracted as movement in mm per second. Immediately after assessment, fish were euthanized using Aqua-Sed™ as per manufacturer’s guidelines.

1.1.7. **Light/Dark Behavioral Analysis**

The raw data files were imported into a custom R script (R Core Team, 2024), available through OSF, where they were standardized against baseline activity levels. This normalization involved calculating each larva's movement as a percentage relative to the mean baseline response. Outliers were identified and excluded when they exceeded the absolute deviation from the median, using the approach described by (Leys et al. (2013) To examine data distribution, residuals from null linear mixed models were visually assessed using QQ-plots and histograms. Since the distribution was non-normal, a zero-inflated gamma (ZIG) generalized linear mixed model (GLMM) was applied, with percentage movement used as the dependent variable. The model accounted for light phase, drug concentration, and time as fixed effects, while Fish ID was included as a random effect. The ZIG-GLMM model was selected to address the presence of 0% movement data and was suitable for analyzing continuous variables with skewed distributions (Zuur & Ieno, 2021). Model fit was assessed using likelihood ratio tests (LRTs), comparing the full model to a null model, with the same process applied to data from the dark phase. Additionally, a Kruskal-Wallis test, followed by Dunn’s post-hoc test, was used to evaluate drug concentration effects on locomotion during the dark phase, with significance levels set at p*** < 0.001, p** < 0.01, and p* < 0.05.

Radar plots and heat maps were generated to illustrate movement in comparison to baseline values. Movements were categorized into darting (>2000%/s), steady swimming (690–2000%/s), slow swimming (0.1–690%/s), or no movement (0%/s). To account for zero responses, a constant of one was added to each observation before calculating percentages. Movement metrics, including total movement, darting, steady swimming, slow swimming, and no movement, were calculated for control groups, with drug treatment groups compared to vehicle controls using log-transformed data (base 10). Distributions of swimming behavior across light and dark phases were evaluated using QQ-plots and histograms, which confirmed non-normal distributions; consequently, Kruskal-Wallis tests and pairwise Wilcoxon-Bonferroni post-hoc tests were applied to analyze these data.

The observed power for each exposure was calculated, with all behavioral tests showing sufficient power (> 0.8 or 80%), indicating no further sample size calculations were necessary.

1.1.8. **Adult Behavioral Analysis**

The raw data files were analyzed by summing the movement per minute per fish in Microsoft Excel (Microsoft Excel, 2024). The movement was then averaged over the 10-minute recording period for each fish and an overall average per treatment group calculated. Using a custom R processing script, the group comparisons were calculated with a one-way analysis of variance (ANOVA) and post-hoc comparisons calculated using Tukey’s multiple comparisons test. A power analysis was also calculated with the coding script.

1.1.9. **Larval Light-only Analysis**

The raw data files were analyzed by summing the movement (distance travelled in mm) per five minutes per fish in Microsoft Excel (Microsoft Excel, 2024). These data were then imported into a custom R coding script and plotted as distance travelled including a 30-minute baseline (pre-exposure) and 80-minute post-exposure locomotor response.

1.1.10. **Light-phase Transition Analysis**

The minute before, during and after a light phase change (5-minutes, 15-minutes and 25-minutes) were analyzed as mean distance travelled per exposure group in GraphPad Prism 10 (GraphPad Prism, Version 10) by conducting a two-way ANOVA comparing the light phase change (5-minutes, 15-minutes and 25-minutes) as well as the pre-, during- and post-light phase change. Significance values were set at: *p***** < 0.0001, *p**** < 0.001, *p*** < 0.01, and *p** < 0.05. Data visualisations were produced in R.

1.1.11. **Light Sheet Microscopy**

Light sheet microscopy was performed using a customized setup inspired by the OpenSPIM platform, as previously detailed by Winter et al. (2017, 2021). Transgenic zebrafish larvae expressing *elavl3:GCaMP6* were pre-treated for 20-minutes with either water, 1 μM fentanyl, 2.0% ethanol or a 1 μM fentanyl and 2.0% ethanol combination in a 24-well plate. The larvae were then transferred to a solution containing the exposure drug concentration plus the neuromuscular blocker tubocurarine (4 mM). After approximately one minute, when muscle tone was lost, larvae were embedded in 1.4% low melting point (LMP) agarose mixed with the treatment solution and drawn into a borosilicate glass capillary (940 μm internal diameter) and sealed with 1.5% LMP agarose. Mounted larvae were oriented head-down and imaged using the light sheet microscope. Sequential horizontal place imaging was performed from the dorsal to ventral surface, covering a total depth of 220 μm across 10 equidistant Z-planes. Each imaging session, lasting around 6-minutes, generated a 200-frame Z-stack. Larvae were confirmed to be viable after imaging through observation of a normal heart rate and blood flow.

The imaging system employed a 288 nm Argon laser (Melles Griot, the Netherlands) for excitation and a 20x 0.5 NA objective lens (Olympus, UK) with a 1x magnification intermediate. Fluorescence emission was collected through 525/50 nm and 495 nm bandpass filters (Chroma, Germany). Image capture was performed with a 5.5 MP Zyla sCMOS camera (Andor, Northern Ireland) operating at 30 frames per second, with a resolution of 640 x 540 pixels, 4 x 4 binning, and 40 ms exposure. Microscope and stage functions, including rotational axis adjustments, were managed using the μManager software integrated with the OpenSPIM plugin.

1.1.12. **Image Processing and Fluorescence Analysis**

Image data were analyzed through a custom-built Python pipeline, incorporating libraries such as Scikit-image, SciPy, and Scikit-Learn, previously described by Winter et al. (2017, 2021). The pipeline began with 3D shift correction and voxel down-sampling (3 × 3 × 1 block averaging) to enhance computational efficiency. Baseline correction was performed using a sliding minimum filter, and this baseline was subtracted from the raw fluorescence data.

All image sets were aligned to the Z-brain Atlas (Randlett et al., 2015) which provided a reference template for 3D image registration. Initially, key points from 14 anatomical landmarks were manually identified for each imaging series on a representative z-stack. Optimal affine transformations were then calculated using the Umeyama algorithm to align these landmarks to their reference counterparts in the reference brain. This method allowed the registration of 45 predefined regions of interest (ROIs) corresponding to major brain structures. Registration accuracy was visually confirmed by overlaying the template with the datasets, and any misaligned images were manually corrected by adjusting the key points. Median alignment error was calculated as the distance between reference and aligned points.

For each ROI, voxel fluorescence intensity data were summarized using the mean, median, and standard deviation, with the median chosen as the primary measure due to its robustness against minor spatial misalignments. Temporal activity profiles were smoothed with a Gaussian filter (sigma = 1.5) to reduce noise. Peaks in fluorescence were identified based on a threshold of two standard deviations above baseline and were considered significant if they persisted for at least two imaging cycles (>1.875 seconds). Metrics such as peak height, width, interval, area under the curve (AUC), and total number of peaks were also calculated for each subject.

Statistical comparisons of fluorescence intensity between fentanyl-exposed and vehicle-treated controls were conducted using unpaired t-tests on the median fluorescence intensity values for each ROI following normality checks. Percent change in fluorescence intensity (%ΔF/F) was calculated using the following formula:

ΔF/F=(F1−F0)/F0×100

Where F1 represents fluorescence intensity in the treatment group, and F0 represents the control group intensity. Visualizations of results, including ROI activity and fluorescence dynamics, were created using GraphPad Prism 10 (GraphPad Prism, Version 10), Microsoft Excel, and custom Python scripts. All code and analysis tools are available on GitLab and OSF for reproducibility.

1.1.13. **Figures Statement**

All figures were compiled and produced in BioRender.com to ensure consistency.

1.2. **Supplementary Materials: Swim Phenotypes**

**Supplementary Table 2** includes an overview of the different swim phenotypes that are seen in the larval light/dark assay and their potential interpretations.

**Supplementary Table 2:** Overview of the swim phenotypes, the movement parameters required compared to control response and interpretation of the response.

| **Phenotype** | **Percentage of control response (%/s)** | **Interpretation** |
| --- | --- | --- |
| Darting | > 2000 | Indicator of initial dark escape/light-seeking response; Anxiety-like response |
| Steady Swim | 690 – 2000 | Continuous swimming commonly seen following drug exposure (not common with controls) |
| Slow Swim | 0.1 – 690 | Any reduced swim response from control |
| Zero Moves | 0 | Freezing; sedation; paralysis; death |
| Total Moves | N/A | Combination of different swim responses showing a complex behavioral response |

1.3. **Supplementary Materials: Figures**

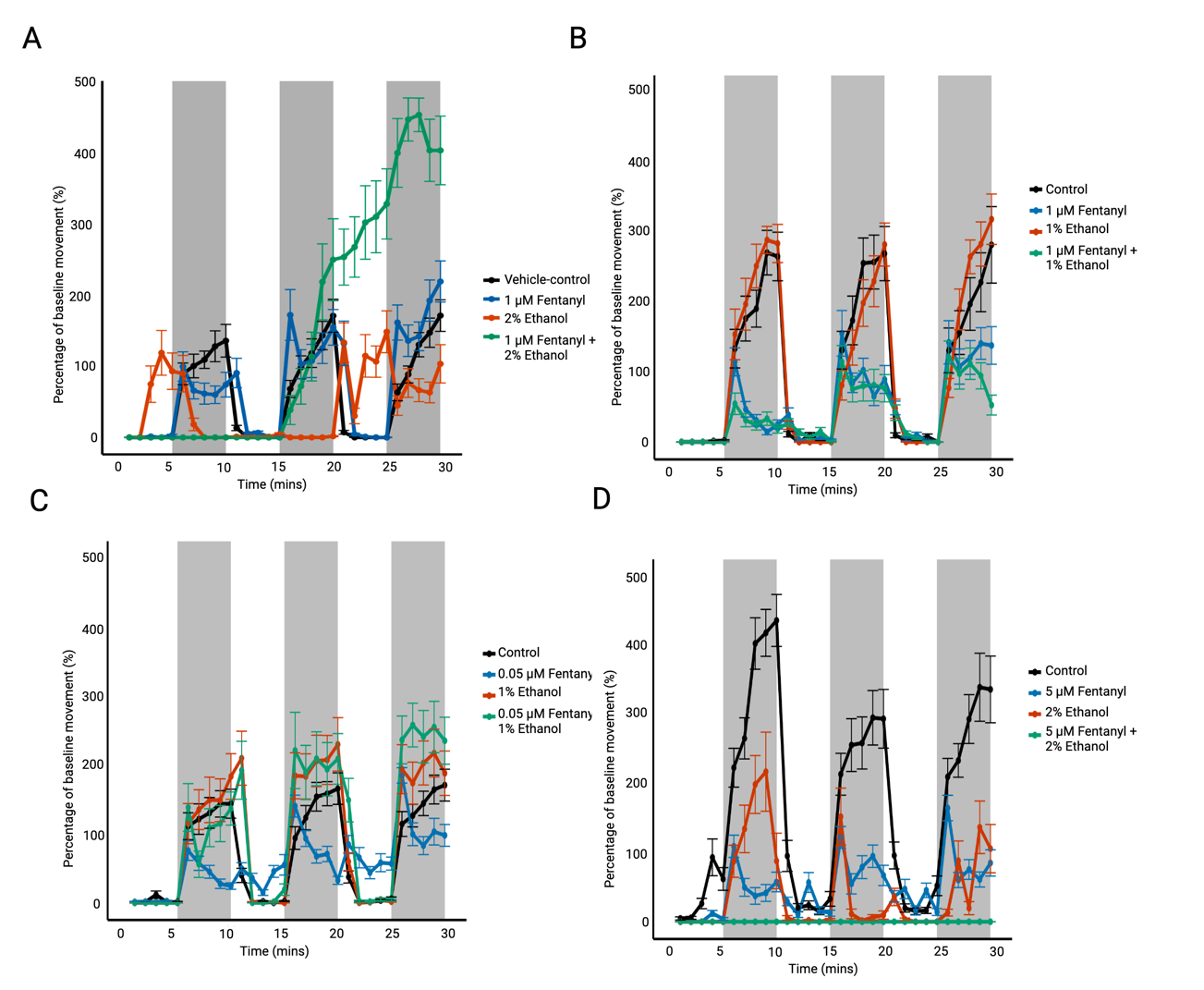

**Supplementary Figure 1: Preliminary 0-30-minutes post-exposure concentration-response poly-drug exposure findings using the larval light/dark assay.** Preliminary light/dark locomotion findings with 4 dpf larval exposed to a range of fentanyl-ethanol poly-drug concentrations from which the 1 μM fentanyl and 2.0% ethanol (A) was identified as warranting further investigation. Data plotted as mean ± SEM per minute. (*n* = 16, preliminary evaluations).

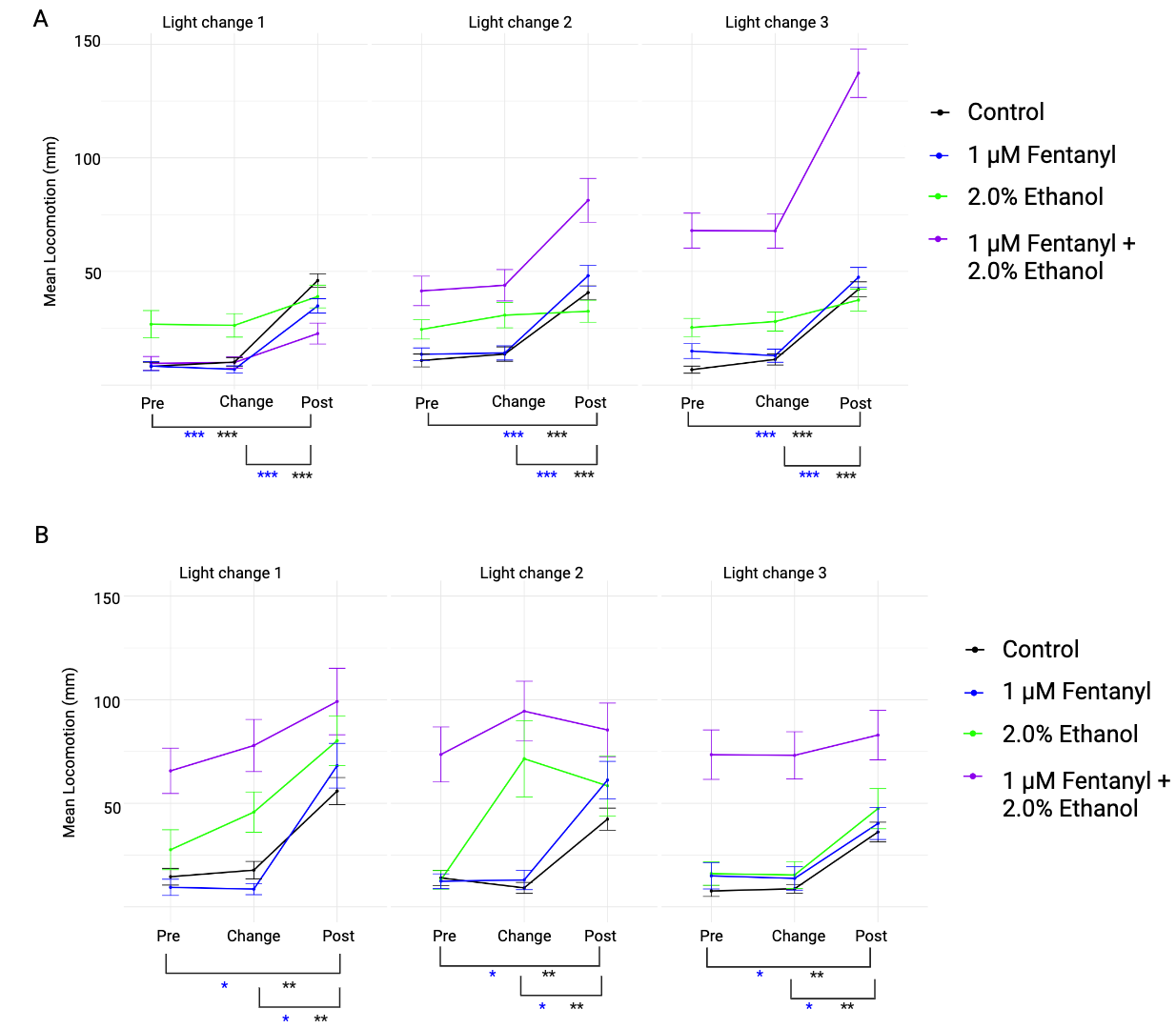

**Supplementary Figure 2: The mean larval locomotion one-minute pre-, during, and post- light phase change for the 5-minute, 15-minute and 25-minute light phase changes.** The average locomotion represented as distance travelled (mm) for larvae during the 5-minute, 15-minute and 25-minute light phase changes for the minute before, during and after the light changes. (**A**) represents the 0-30-minute data and (**B**) the 60-90-minute data. (*n* = 98, observed power = 0.997).

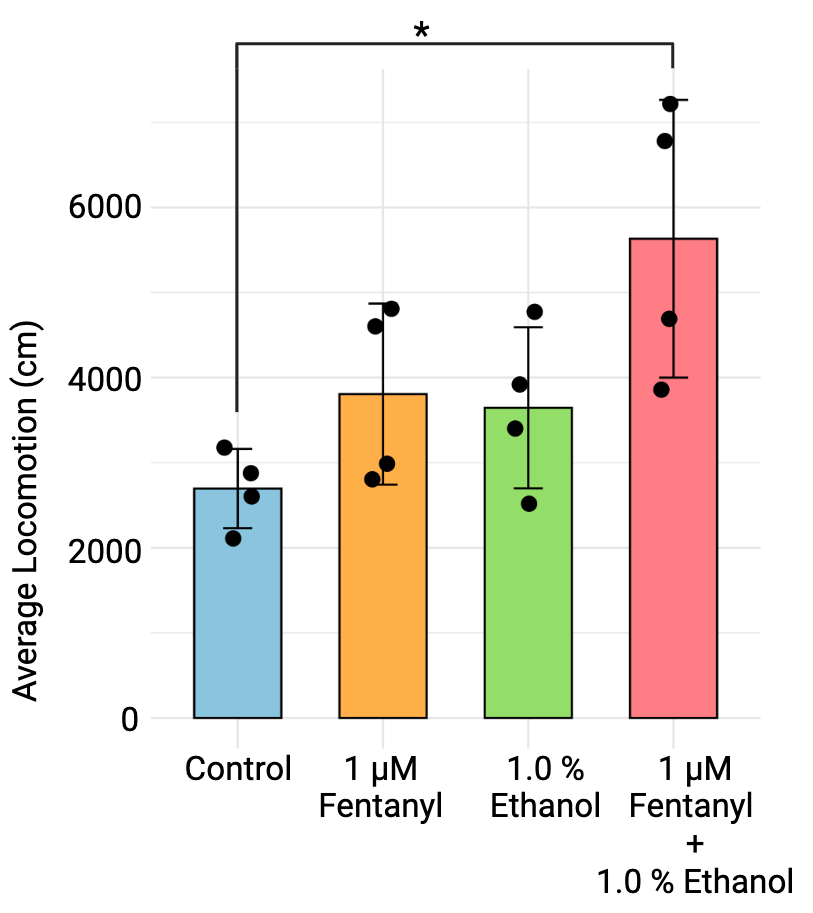

**Supplementary Figure 3: Confirmation of hyperactive behavioral phenotype in adult zebrafish exposed to the fentanyl-ethanol combination.** The average distance traveled (cm) over a 10-minute recording period for 8-month post-fertilization AB adult zebrafish exposed to water, 1 μM fentanyl, 1.0% ethanol or a fentanyl-ethanol combination. Data plotted as mean ± SEM. (*n* = 4, observed power > 0.8).

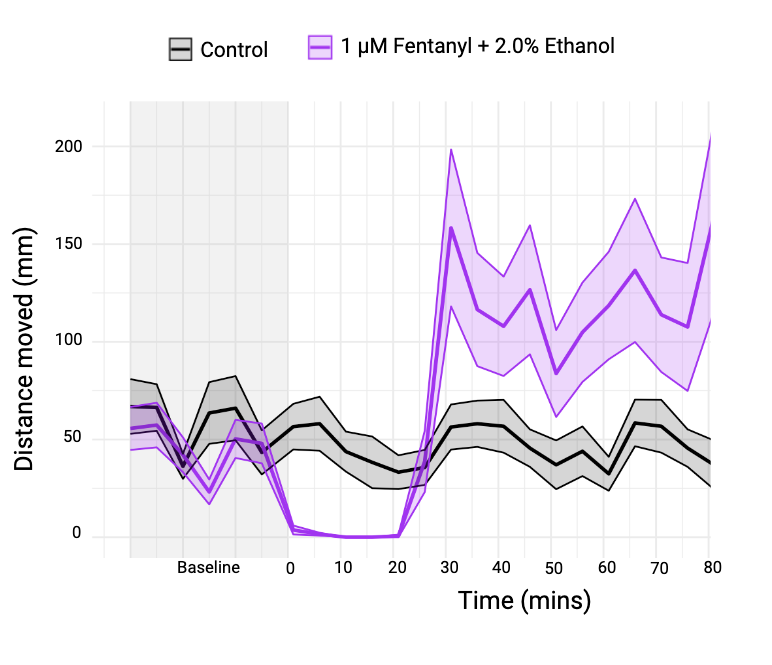

**Supplementary Figure 4: Light-only larval zebrafish locomotor response following administration of 1 μM fentanyl and 2.0% ethanol.** Analysis represented as mean distance travelled (mm) over the 80-minutes recording period following exposure to the fentanyl-ethanol combination including a 30-minute baseline recording (pre-exposure) for reference. Data presented as mean (solid line) ± SEM (shaded area). Observed power = 0.997.

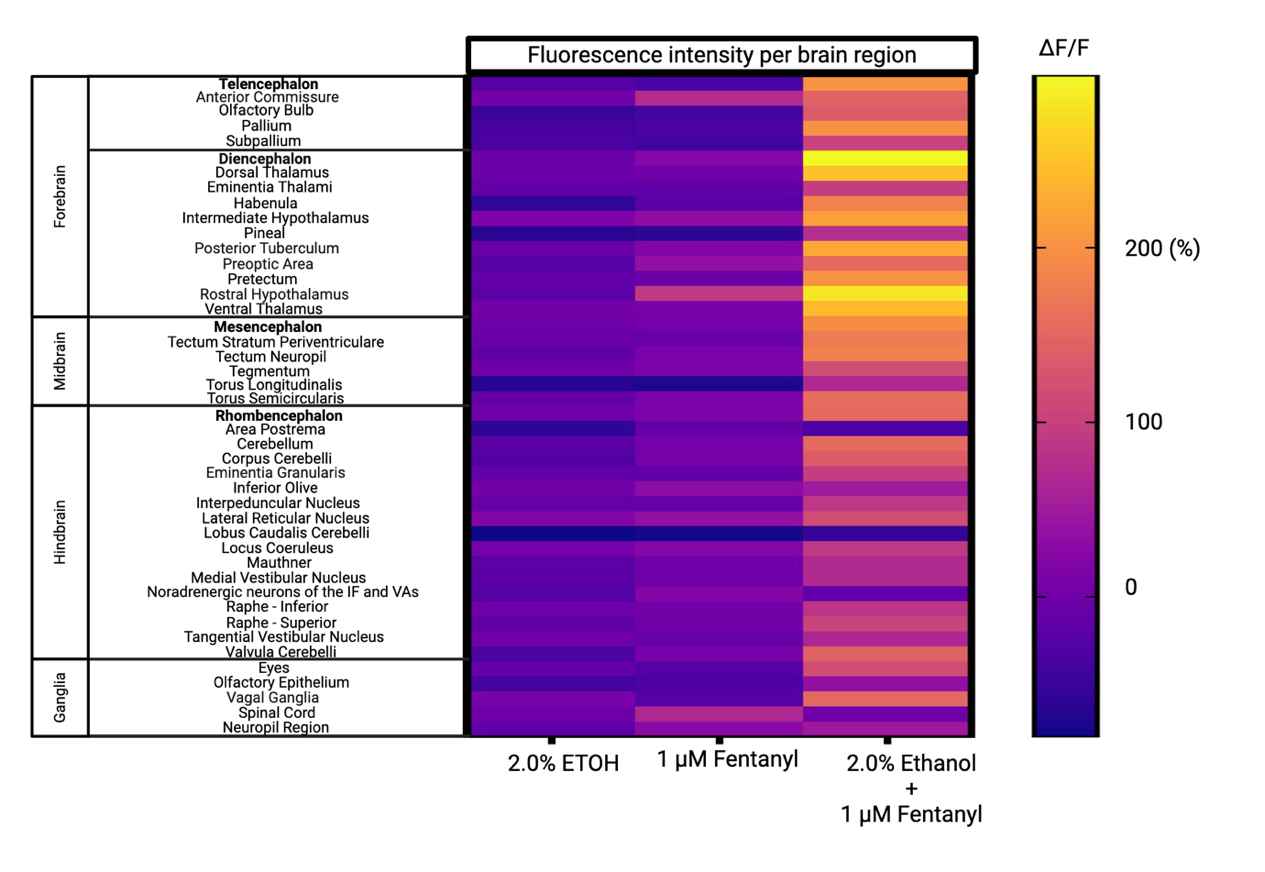

**Supplementary Figure 5:** **The percentage change of fluorescence intensity per brain region.** The percentage change in fluorescence intensity of 2.0% ethanol, 1 μM fentanyl and the fentanyl-ethanol combination compared to vehicle-controls presented as (Δf/f = (f_1_ – f_0_)/f_o_ *100, where f_1_ = peak fluorescence intensity and f_0_ = control fluorescence intensity within each region of interest (ROI).

**Supplementary Materials: Videos**

**Supplementary Video 1:** **Video recording of 4 dpf zebrafish larvae exposed to the fentanyl-ethanol combination.**

**[video attached separately]**

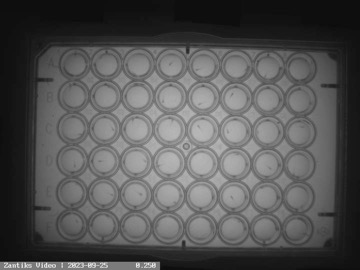

Video recording from the Zantiks MWP behavioural unit for the light only recording of the response. Row A: controls, B: 1 μM fentanyl, C: 2.0 % ethanol, D, E, F: 1 μM fentanyl-2.0 % ethanol combination. The video is sped up ~25x to account for the length of recording.

**Supplementary Video 2: Video recording obtained from the Zantiks AD behavioral unit of an adult zebrafish exposed to the fentanyl-ethanol combination.**

**[Video attached separately]**

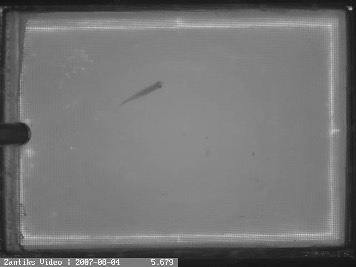

The 10-minute live behavioral recording of an adult AB zebrafish exposed to the fentanyl-ethanol combination. The video has been sped up 30x.
